# Ecological opportunity and evolutionary relatedness predict parasite sharing differently across mammalian taxonomic scales

**DOI:** 10.64898/2026.11.08.765232

**Authors:** Nicole Nova, Erin A. Mordecai

## Abstract

As global change brings species into new contact, understanding the ecological and evolutionary drivers of parasite sharing is essential for anticipating and mitigating disease emergence. Yet, the relative importance of geographic proximity and evolutionary relatedness among host species in predicting parasite sharing remains unresolved. We tested whether their relative importance changes across host taxonomic scale using 30,310 recorded associations between 680 mammalian and 11,107 parasite species. Geography accounted for 86.6% importance within genera, declining to 44.7% across all mammals, shifting toward evolutionary relatedness predominating. Parasite sharing increased nonlinearly with geographic proximity and host relatedness, and the overall trend varied among parasite types. Notably, accounting for host taxonomic scale improved parasite-sharing prediction accuracy. To explore the consequences of biotic homogenization, we applied a counterfactual scenario of reducing geographic distance by 99%, which increased mean predicted parasite sharing to 13.71% within genera and 1.50% at the class level. Together, these findings suggest that ecological opportunity is especially consequential among close relatives whereas evolutionary similarity is increasingly important among geographically distant species. Accounting for evolutionary scale can therefore improve sharing predictions and help identify which new host encounters warrant surveillance for cross-species transmission and disease emergence.

## Introduction

Understanding why host species share parasites is central to anticipating disease emergence and protecting human and wildlife health^1^. Cross-species transmission depends on opportunities for exposure and the ability of a parasite to infect a recipient host^2,3^. Patterns of parasite sharing, defined as infection by the same parasite species in different host species, can therefore reveal ecological and evolutionary conditions associated with cross-species transmission^1,4,5^. Such events can have profound consequences: the emergence of HIV followed transmission from nonhuman primates to humans, while canine distemper outbreaks have spread beyond domestic dogs and threatened wild carnivores and other species^6,7^. Changes in climate and land use are bringing host species into new geographic settings and altering their opportunities for contact, increasing the need to understand which host encounters are most likely to permit parasite sharing^8–10^. Identifying the conditions that allow parasites to cross species boundaries can inform surveillance and interventions that address disease emergence before outbreaks occur^1,11^.

Previous studies have repeatedly identified geographic proximity and evolutionary relatedness as major predictors of parasite sharing among mammals^12–16^. Species with overlapping ranges may encounter the same parasites through direct contact, shared environments, vectors, or intermediate host species, making geography a useful predictor of opportunities for exposure^15–17^. Closely related species may share physiological and immunological characteristics that allow the same parasites to infect them, making phylogenetic distance a predictor of host compatibility^2,3,18^. Associations with geography and relatedness have been documented in primates, carnivores, ungulates, bats, and rodents, as well as in analyses spanning mammals^12,14–16,19^. Ecological and life-history traits also contribute to parasite sharing, with overall trait similarity predicting parasite assemblage similarity in carnivores and habitat use, longevity, and reproductive traits associated with species’ roles in mammalian parasite-sharing networks^14,20^. Recent studies of bat viromes, rodent ectoparasites, and mammalian mites further demonstrate the value of considering host geography and evolutionary history together^21–23^. These predictors offer a basis for identifying potential cross-species infections as host distributions and opportunities for exposure change^1,5,8,10^.

Despite agreement that geography and evolutionary relatedness predict parasite sharing, studies have reached different conclusions about their relative importance^12,14–16^. In primates, evolutionary relatedness was the stronger predictor across pathogens overall, whereas geography was stronger for viral sharing, and geographic overlap also outweighed phylogenetic similarity in the viral-sharing analysis of ungulates^12,15,16^. In contrast, Albery and colleagues found that phylogeny explained more variation than geography in viral sharing across mammals, and suggested that the broader evolutionary range of host species could explain the difference from narrower comparisons^16^. Previous analyses have varied in their host groups, parasite types, sampling coverage, and statistical methods, making comparisons among studies insufficient to determine whether taxonomic scale explains these contrasting rankings^13,15,16,19^. A relationship estimated among close relatives may not describe sharing between deeply divergent host species, and models of geographic range shifts depend on how those relationships are specified^8,16^. A direct comparison across taxonomic scales is therefore needed to distinguish a change in predictor importance from differences among separately assembled studies^16,24^.

Here, we tested whether the relative importance of geography and evolutionary relatedness changes with taxonomic scale using parasite records for 680 mammalian species from 13 orders. We compared pairs within genera, families, and orders, and across the mammalian class, using a modeling framework that accommodates nonlinear relationships and accounts for study effort. We hypothesized that geographic proximity is the stronger predictor among close relatives, while evolutionary relatedness becomes more important as comparisons include more divergent species. We examined whether this pattern extends across major parasite types and how additional host traits contributed at each scale. We then assessed whether accounting for taxonomic scale improves predictions of parasite sharing and whether models trained on close relatives can predict sharing among distant relatives, and vice versa. Finally, we evaluated how predicted parasite sharing, while accounting for taxonomic scale dependence, changes under potential scenarios where geographic separation is reduced, for example through biotic homogenization of mammalian communities. Together, these analyses tested whether a crossover in geographic and evolutionary importance across taxonomic scales helps reconcile previous findings and changes predictions of which novel host encounters are most likely to permit parasite sharing.

## Results

Our dataset contained 30,310 distinct host–parasite associations connecting 680 placental mammal species from 13 orders to 11,107 recorded parasite taxonomic names. Host species were retained if they had at least ten recorded parasite names and complete geographic, evolutionary, taxonomic, and study-effort information. The resulting network connected 68 of the 78 possible pairs of mammalian orders through at least one shared parasite name (Fig. 1). Bats, rodents, carnivores, even-toed ungulates, and primates contained the greatest numbers of included host species and recorded parasite names. The five strongest sharing connections, measured by shared-parasite counts, were between even-toed ungulates and carnivores, carnivores and rodents, even-toed ungulates and rodents, carnivores and primates, and even-toed ungulates and primates. Carnivora had the greatest total shared-parasite count across its connections. Artiodactyla (even-toed ungulates), Carnivora (carnivores), Chiroptera (bats), Eulipotyphla (insectivores), Rodentia (rodents), and Primates had the most connections to other orders. Recorded sharing therefore extended widely across mammalian lineages, with several species-rich orders forming highly connected parts of the network. The evolutionary relationships among the included host species are shown in Fig. S1.

**Fig. 1.**
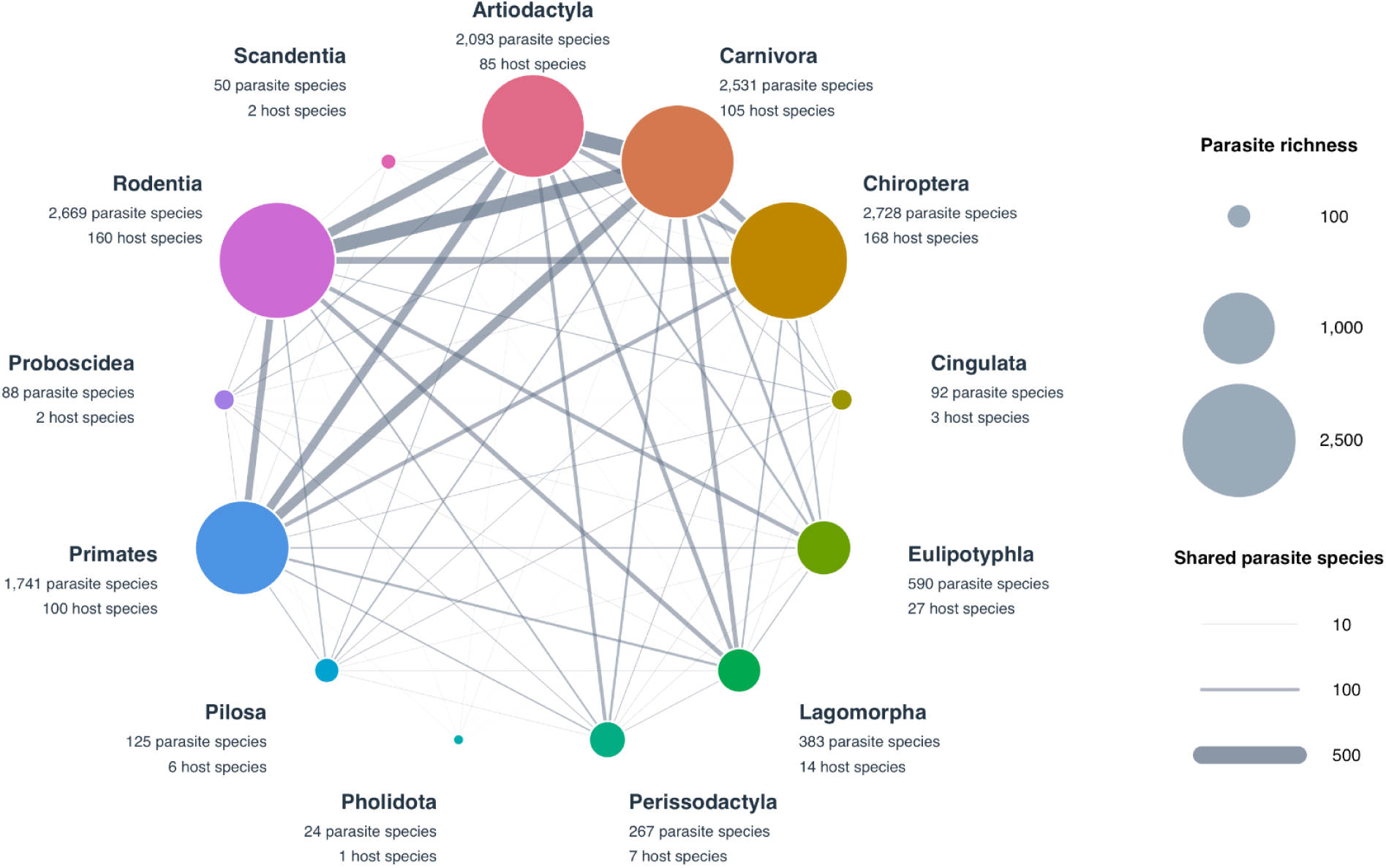
Recorded parasite sharing among mammalian orders in our dataset. Nodes represent the 13 mammalian orders, with colors identifying orders and node area proportional to parasite richness across their included host species. Labels give parasite richness and the number of host species. Edges connect orders with at least one shared parasite species. Edge width is proportional to the shared count. All 68 nonzero overlaps among 78 possible order pairs are shown. Circular positions are alphabetical and do not encode geographic or phylogenetic distance. A parasite recorded in multiple orders contributes to each applicable order-pair overlap, so edge counts are not additive unique totals. The dataset contains 680 host species, 11,107 distinct recorded parasite taxonomic names, and 30,310 distinct host–parasite associations.

### Geography is more important among close relatives, while evolutionary relatedness becomes more important across mammals

We first compared models fitted to increasingly broad sets of host pairs, encompassing species within the same genus, family, or order and any pair in the mammalian class. All models included geographic distance, evolutionary distance, and differences in study effort as predictors. The response variable was the proportion of parasite community overlap between species^12,25^. We measured the relative importance of geography and evolutionary relatedness by their respective contributions to the fitted effects.

Geographic importance declined as the comparison broadened from genus to class (Fig. 2A). It accounted for 86.6% of combined geographic and evolutionary importance within genera (95% interval: 66.4–97.9%), 56.3% within families (45.2–71.2%), 49.2% within orders (43.9–54.5%), and 44.7% across mammals (40.3–48.6%). Evolutionary relatedness therefore accounted for 55.3% at the broadest scale (51.4–59.7%). Bootstrap comparisons supported greater geographic importance within genera (*P* < 0.001) and greater evolutionary importance across mammals (*P* = 0.018), but did not distinguish their contributions within families (*P* = 0.374) or orders (*P* = 0.759). Together, these results show a shift from geographic proximity as the stronger predictor of parasite sharing among close relatives to evolutionary relatedness as the stronger predictor across mammals.

**Fig. 2.**
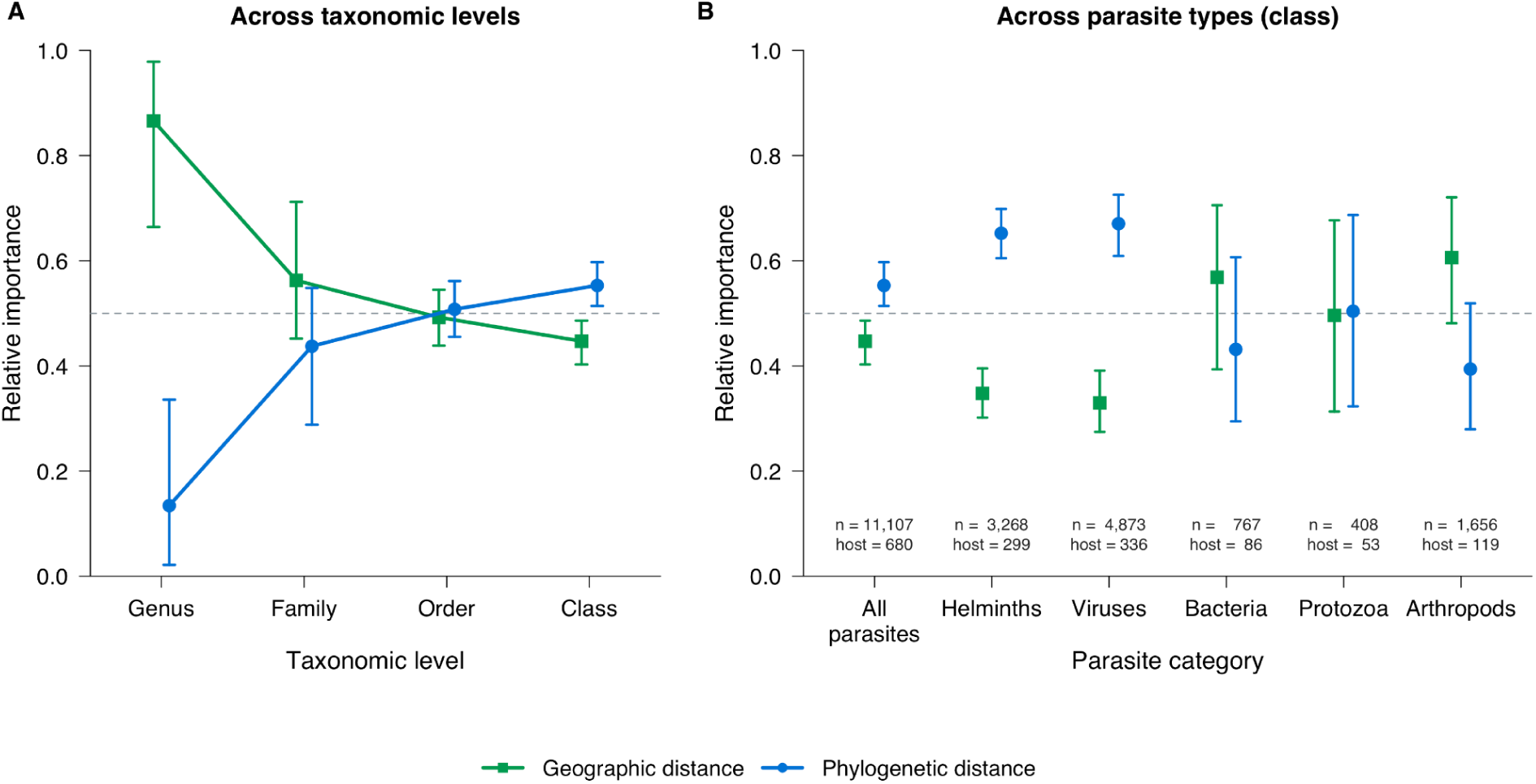
Relative importance of geographic distance and evolutionary relatedness in predicting parasite sharing among mammalian host species. (A) Geographic and phylogenetic importance across host pairs belonging to the same genus, family or order, and across all mammalian hosts at the class level. (B) Relative importance across all parasites and five parasite types at the class level. The number of host species for each parasite type is added after “host” and “n” indicates the number of parasite species included. Type-specific models include hosts with at least 10 recorded parasite taxa of the corresponding type. Green squares represent geographic distance between host range centers, and blue circles represent phylogenetic distance between host species. Relative importance is each distance predictor’s summed fitted spline coefficients (from the regression model) divided by the combined geographic and phylogenetic coefficient sums. The two shares sum to one, and the dashed line marks equal importance. Points show fitted estimates and error bars show 95% percentile intervals from 1,000 host-bootstrap replicates. Parasite sharing refers to similarity between recorded parasite communities.

Further, the fitted geographic and evolutionary distance relationships with parasite sharing were nonlinear (Fig. 3). Parasite community dissimilarity increased steeply over short geographic distances before approaching a plateau, and the curves were similar at family, order, and class levels, with a somewhat different shape within genera (Fig. 3A). Evolutionary distance responses differed more strongly among taxonomic scales (Fig. 3B). Parasite community dissimilarity increased rapidly before leveling off within genera, continued rising within families, and showed an initial rise followed by a plateau and a renewed increase within orders. Across mammals, the relationship was S-shaped, with a gradual initial increase, a steeper rise at intermediate evolutionary distances, and leveling off at the greatest distances. Thus, as the taxonomic scale broadened, the relationship between parasite sharing and evolutionary distance changed more than the relationship between parasite sharing and geographic distance.

**Fig. 3.**
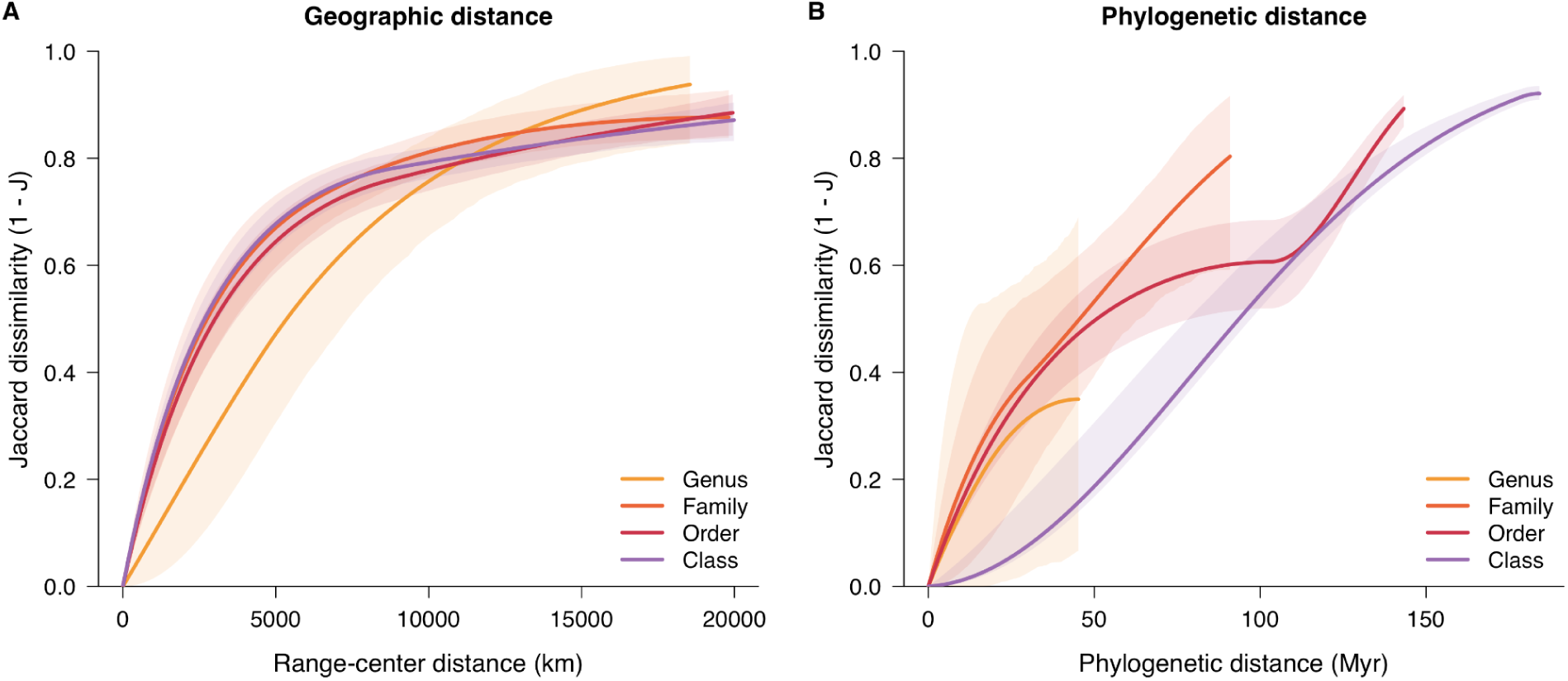
Geographic and evolutionary distance effects on parasite community dissimilarity across mammalian taxonomic scales. Curves show the isolated effects of (A) geographic distance between host species’ range centers and (B) phylogenetic distance measured as the sum of evolutionary branch lengths separating two host species in millions of years (Myr). Colors denote models fitted to host pairs within the same genus, family, order, or class. Lines show fitted effects and shaded bands show pointwise 95% percentile intervals from 1,000 host-bootstrap replicates. Larger values indicate greater parasite community dissimilarity (less sharing).

Separate analyses showed that the relative importance of geography and evolutionary relatedness varied among parasite types. At the class level, evolutionary relatedness was more important than geographic distance for helminths and viruses, but the two were of similar importance for bacteria, protozoa, and arthropod parasites (Fig. 2B; Fig. S2). Across all mammals, evolutionary relatedness accounted for 65.2% of the importance for helminths (95% interval: 60.5–69.9%; 299 host species) and 67.1% for viruses (60.9–72.5%; 336 host species). Corresponding estimates were 43.2% for bacteria (95% interval: 29.4–60.7%; 86 host species), 50.4% for protozoa (32.3–68.7%; 53 host species), and 39.4% for arthropods (27.9–51.9%; 119 host species), but the overlapping intervals for these three types suggested equal geographic and evolutionary importance (Fig. 2B). Across parasite types, geographic importance generally decreased and evolutionary importance increased as taxonomic scope broadened, although individual steps were not always monotonic (Fig. S2). Bacteria and arthropods retained higher geographic point estimates even across mammals. The crossover between geographic and evolutionary importance therefore differed among parasite types, and was not evident within the examined scales for every type, but the overall trend of decreased geographic importance relative to evolutionary relatedness as taxonomic scale broadened remained.

We further examined whether differences in host range breadth of a parasite could explain variation in the crossover between geographic and evolutionary importance among parasite types. Indeed, recorded host ranges varied among parasite types (Fig. S3). The proportion of parasite names recorded in only one retained host species ranged from 53.4% for bacteria to 80.0% for viruses (Fig. S3A). Bacteria tended to have broader host ranges and viruses narrower host ranges than other parasite types (Fig. S3B). However, a virus (rabies lyssavirus) also had the maximum recorded host range of 165 host species (Fig. S3B). Thus, bacteria may infect a broader range of host species, with geography remaining more important at broader taxonomic scales, whereas viruses may be more specialized, with some exceptions, and more constrained by evolutionary relatedness. Other parasite types may fall between these extremes. However, these estimates describe the compiled records, rather than establishing biological specialization or the complete host range of each parasite. Whether parasites with broader biological host ranges show a crossover at greater evolutionary distances remains a question for comparisons extending beyond mammals.

Moreover, adding one host trait at a time to the models revealed other predictors of parasite sharing at certain taxonomic scales (Fig. S4). Body mass, body length, and maximum longevity generally contributed more at the class, family, and order levels than within genera, although intervals overlapped and did not establish a consistent increase across all four scales (Fig. S4A). Conversely, habitat breadth contributed most within genera, families, and orders, on average and less across mammals (Fig. S4B), while litter size and population density contributed at all scales (Fig. S4C). Home range size, activity cycle, terrestriality, and trophic level generally made weaker contributions to the fitted effects (Fig. S4D). Together, these results suggest that host traits provide additional information about parasite sharing, but their contributions depend on the trait and taxonomic scale.

### Taxonomic scale affects predicted parasite sharing among host species

To test the importance of accounting for taxonomic scale in parasite sharing prediction, we applied the scale-dependent models fitted to the genus and class levels to species pairs within genera and to more distant host species pairs at the mammalian class level. We evaluated both models on the same held-out species pairs, with entire genera withheld during training to limit information shared between training and evaluation sets.

Among pairs within genera, the genus-trained model had a root mean squared error of 6.93 percentage points of sharing, compared with 17.34 percentage points for the model trained at the class level (Fig. 4). These percentage points express an absolute difference between percentages, since the errors use the percentage scale of parasite sharing. To compare performance across taxonomic scales, we also divided each model’s error by the error of a baseline that assigned every test pair the mean sharing observed among the corresponding training pairs. These standardized errors are unitless ratios, with values below one indicating better performance than this baseline. Models trained on within-genus pairs predicted within-genus sharing relatively well, with a standardized error of 0.83 (95% interval: 0.76–0.93), whereas the class-trained model performed poorly for these pairs, with a standardized error of 2.09 (1.73–2.58) (Fig. 4A). The reverse pattern occurred for pairs at the class level: the class-trained model performed well, with an error of 0.99 percentage points and a standardized error of 0.96 (0.94–0.98), whereas the genus-trained model performed poorly, with an error of 4.43 percentage points and a standardized error of 4.28 (3.77–4.85) (Fig. 4B). Thus, models predicted parasite sharing more accurately when trained and evaluated at the same taxonomic scale, indicating that accounting for taxonomic scale and the relative contributions of geography and evolutionary relatedness is important for prediction.

**Fig. 4.**
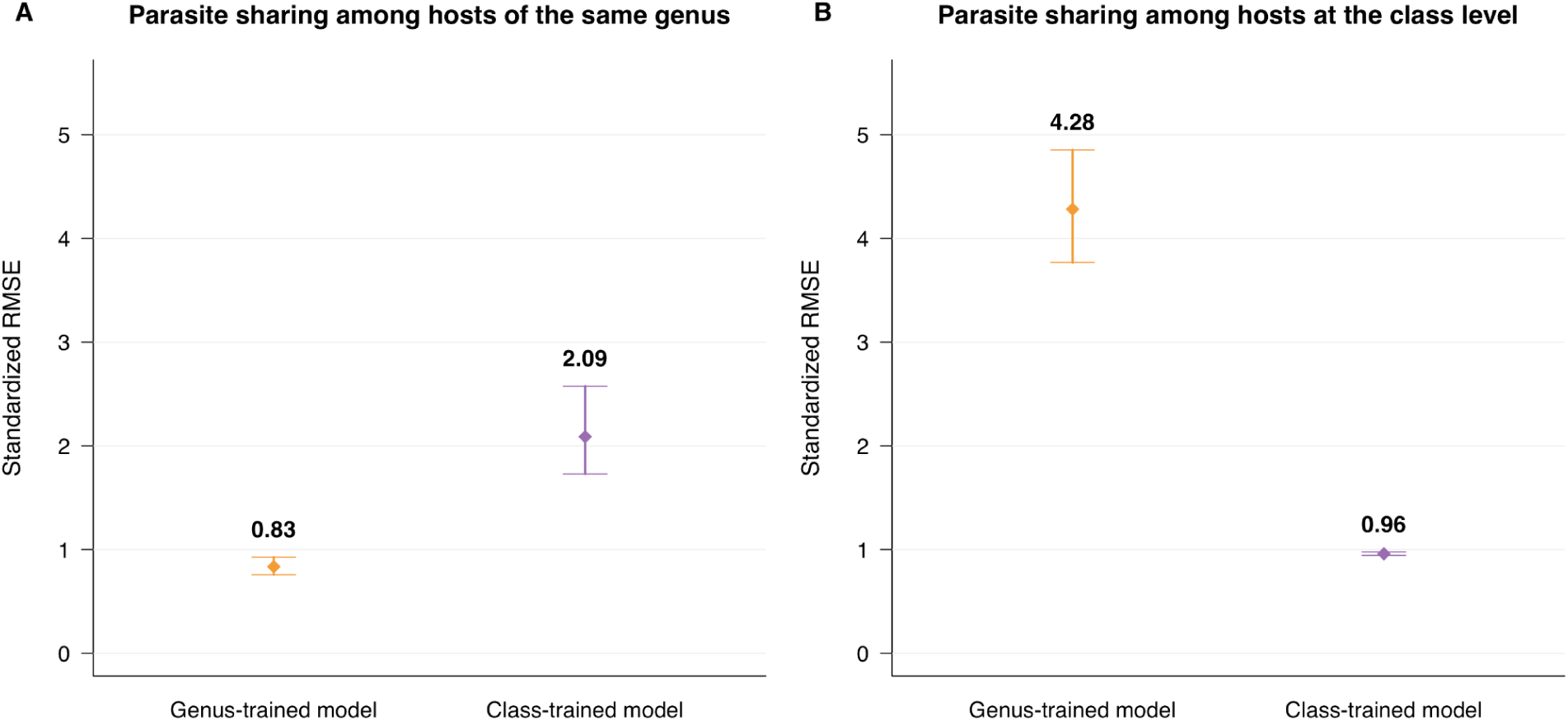
Predictive performance of genus-trained and class-trained models across taxonomic domains. In the genus-trained model, geographic distance is more important than phylogenetic distance between host species. Conversely, in the class-trained model, phylogenetic distance is more important than geographic distance between host species (Fig. 2). Models trained on host pairs from the same genus (yellow) or at the class level (purple) predict recorded parasite sharing among (A) hosts of the same genus and (B) hosts at the class level. Both models include geographic distance, phylogenetic distance and publication counts to account for study effort. Performance is evaluated using five-fold whole-genus holdout. All species in a test genus are excluded from both models’ training pairs. Points show standardized root mean squared error (RMSE), calculated as model RMSE divided by the RMSE of a constant predictor using the corresponding target’s training-data mean sharing. Error bars are approximate 95% percentile intervals from 2,000 paired genus-block resamples.

### Reduced geographic separation predicts greater parasite sharing among closely related host species

We then assessed how reduced geographic separation could affect parasite sharing as global change drives species movements, including potential climate- and land use-driven range shifts and biotic homogenization driven by global trade and travel. To do so, we used the scale-dependent models to counterfactually reduce the distance between host species’ range centers while holding evolutionary distance and study effort fixed.

Before any geographic distance reductions, the observed mean parasite sharing was 7.85% within genera, 4.75% at the family level, 1.97% at the order level, and 0.42% at the mammal class level (Fig. 5A). Under a 99% reduction in geographic distance, mean predicted sharing reached 13.71%, 12.93%, 6.42%, and 1.50%, respectively (Fig. 5A). Increases relative to each model’s fitted baseline were 5.86 percentage points within genera (95% interval: 3.41–10.10), 8.11 at the family scale (6.55–9.84), 4.42 at the order scale (3.75–5.13), and 1.07 at the class scale (0.90–1.23) (Fig. 5B). A smaller, 25% distance reduction yielded increases of 1.09, 0.78, 0.36, and 0.07 percentage points, respectively (Fig. 5B). Close relatives had the highest final predicted sharing, but the largest increase under the 99% reduction occurred at the family level. These results suggest that reducing geographic barriers can substantially increase sharing among close relatives, while predicted sharing remains lower among distant relatives. The magnitude of the increase nevertheless depends on initial separation and the fitted response at each scale.

**Fig. 5.**
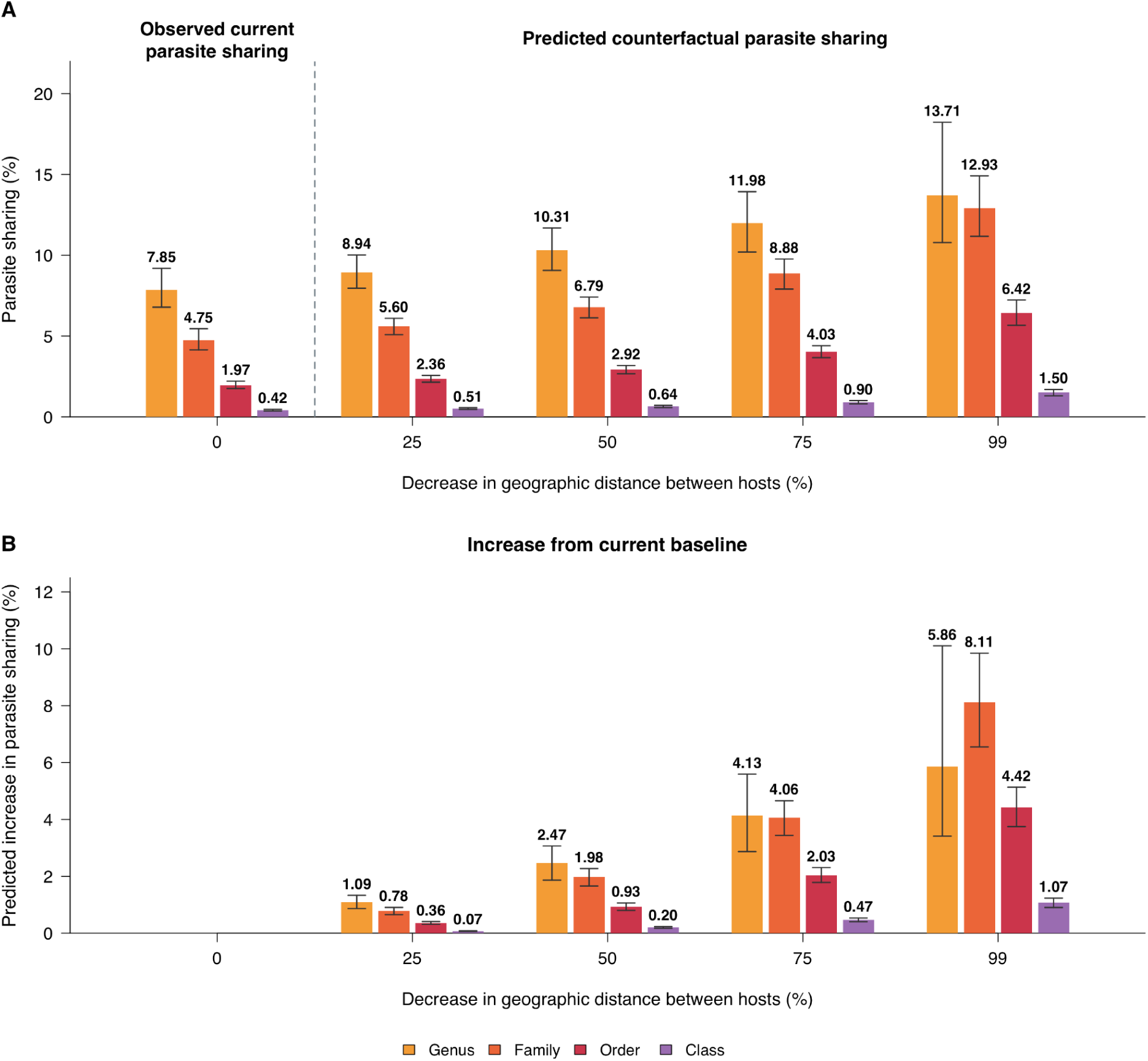
Observed and predicted parasite sharing under counterfactual geographic homogenization scenarios. (A) Mean observed sharing under current conditions (0% distance decrease, left of the dashed separator) and predicted sharing after geographic range-center distances decrease by 25%, 50%, 75% or 99% (right) for the different taxonomic levels: genus (yellow), family (orange), order (red), and class (purple). (B) Predicted increases from the current baseline under the same scenarios for the different taxonomic levels. Parasite sharing is the mean pairwise Jaccard similarity between recorded parasite communities. Predictions use models fitted at the corresponding taxonomic level with geographic distance, phylogenetic distance and publication counts. Phylogeny and study effort remain fixed during the scenarios. Error bars show 95% intervals, where observed means use 1,000 host resamples, whereas future means and within-model baseline-to-future differences use 1,000 compatible host-bootstrap fits. In panel B, increases are absolute percentage-point changes in predicted similarity, not proportional percentage increases. They are calculated from each model’s fitted baseline, rather than by subtracting the observed bars in panel A. Geographic range-center proximity is a contact proxy.

## Discussion

The relative importance of geographic distance and evolutionary relatedness changed systematically with the taxonomic scope of mammalian parasite-sharing comparisons. Geography contributed most strongly within genera, whereas evolutionary relatedness became the stronger predictor across mammals. Both relationships were nonlinear, with greater variation among scales in the evolutionary response than in the geographic response. The balance differed among parasite types: helminths and viruses showed stronger evolutionary contributions across mammals, while the other types had less clearly separated effects. Taxonomic scale affected parasite sharing predictions, and hypothetically reducing geographic separation differentially increased predicted parasite sharing at different scales. Together, these findings help reconcile contrasting importance of geography and phylogeny in previous studies and show why taxonomic scale affects predicted parasite sharing as global change alters species distributions^8,12,15,16^.

A possible mechanism underlying our findings is a scale-dependent balance between opportunities for exposure and compatibility with infection. Close relatives may share physiological and immunological characteristics that allow the same parasites to infect them, making exposure comparatively important^2,3,13^. Among distant relatives, differences in host biology may limit infection or establishment even when geographic barriers diminish^3,18^. Range shifts and species movements may therefore increase parasite sharing particularly among close relatives, while evolutionary differences may continue to constrain sharing among distant relatives. A parasite’s host range may help explain how strongly compatibility constrains sharing, because its ability to infect evolutionarily diverse species may indicate fewer barriers to establishment in distant relatives^26^. Combining this information with scale-dependent sharing models and measurements of host abundance, habitat use, and potential vectors could help identify where surveillance and interventions would be most useful^1,11,27,28^.

The overall shift toward lower geographic and greater evolutionary importance at higher taxonomic levels extended across the parasite types, but the magnitude of this shift and the scale of the crossover differed. Helminths and viruses showed stronger evolutionary contributions across mammals, whereas bacteria, protozoa, and arthropods did not show a clear separation between the predictors at that scale. Differences in infection mechanisms, life cycles, transmission routes, environmental persistence, vectors, and intermediate host species could alter how geography relates to exposure and how host relatedness relates to infection^2,17,26,29^. Recent analyses of rodent ectoparasites, mammalian mites, and regional ectoparasite assemblages likewise show the value of evaluating these relationships in distinct parasite systems^22,23,30^. Our parasite-type analyses do not directly compare transmission strategies, and unequal host coverage limits attribution of the contrasts to parasite biology. Future studies could test whether the crossover occurs at greater evolutionary distances for generalist parasites, using independently characterized host ranges and comparable sampling across parasite types^23,26^.

Host size and life history may further shape sharing through exposure and susceptibility^14,20,31–33^. Body length, mass, and longevity generally contributed more at broader taxonomic scales than within genera, although overlapping uncertainty intervals prevented us from establishing a consistent increase across all scales. Similar body size can link species through predation, competition, and habitat use, creating opportunities for shared exposure even among distant relatives^34–36^. Large terrestrial carnivores, for example, tend to consume larger prey, and predator–prey links can connect ungulates and carnivores that share parasites^15,36^. Body size also scales with metabolic and other physiological processes that could affect the conditions parasites encounter within a host^31,33^. Similar body sizes have evolved across mammalian lineages and environments, potentially allowing ecological or physiological similarities to extend beyond close relatives^37,38^. Similar lifespans may also provide comparable opportunities for exposure and persistence^39^. Experiments or longitudinal observations would help distinguish exposure through shared ecological conditions from similarity in physiological susceptibility^18,40^. Together, these findings suggest that host traits can refine parasite-sharing predictions, but their contribution should be evaluated at the taxonomic scale of the host comparison.

Several data limitations should be considered when interpreting our findings. Geographic and evolutionary distances serve as proxies for exposure and compatibility rather than direct measurements of these mechanisms^3,13,15,16^. Range-center distances omit local contact, seasonal movements, and ecological interactions such as competition and predation, which can connect even distant relatives^15,34–36^. Although ecological networks can help describe these connections, their links indicate potential pathways rather than observed transmission^3,4,41^. Incomplete host–parasite records further constrain interpretation, since study-effort adjustment and minimum-record thresholds cannot establish biological absence from missing records^15,42^, and shared parasite names need not represent genetically identical infections^3,43^. Methods for predicting missing associations can guide targeted sampling^44^, while standardized surveys and finer parasite identification would strengthen comparisons across scales. Our restriction to placental mammals also limits the generality of the findings. Future studies should integrate direct measures of contact and susceptibility with more complete parasite records across marsupials, monotremes, birds, and other host groups to test the mechanisms underlying the crossover from geographic to evolutionary importance and determine how broadly it applies.

Global change is reshaping encounters among host species, and our findings show that their potential impacts on parasite sharing depends on taxonomic scale. Geographic proximity is especially informative among close relatives, whereas evolutionary relatedness becomes more important across deeper host divergences within the Mammal class. Accounting for this shift improves prediction accuracy and changes expectations of sharing as geographic barriers diminish. Surveillance of shifting ranges and newly assembled communities should therefore consider host relatedness alongside opportunities for exposure. Incorporating taxonomic scale into disease risk assessment can help identify host encounters that warrant targeted surveillance and direct limited resources toward opportunities to detect and prevent emerging infections in people, domestic animals, and wildlife.

## Methods

### Data collection and processing

We compiled associations from the Global Mammal Parasite Database version 2^45^, the compilation used by Winternitz and colleagues^46^, CLOVER^43^, VIRION^47^, the Natural History Museum Host–Parasite Database^48^, the mite dataset of Klimov and He^23^, the Brazilian Mammal Parasite Occurrence dataset^49^, and a small resolved residual set of EID2 records^50^. We standardized host and parasite names, reconciled unambiguous synonyms, assigned hosts to genera, families, and orders using the Mammal Diversity Database^51^, and removed duplicate host–parasite associations while retaining information on their sources. The resulting dataset contained wildlife placental mammals.

We measured parasite sharing using Jaccard similarity, J = *s*/(*s* + *a* + *b*), where *s* is the number of parasite names recorded in both species, *a* is the number recorded only in the first species, and *b* is the number recorded only in the second species^12,25^. This measures the shared fraction of the combined recorded parasite community, with values from zero (no sharing) to one (identical recorded communities). For modeling, we used dissimilarity, 1 − J. A shared parasite name indicates a documented association in both species.

Data on morphological and ecological factors, including host life history traits and coordinates of geographic ranges (Table S1), were obtained from the publicly available PanTHERIA database^52^. Geographic distance was the distance in kilometers between host species’ range centers. Centers were calculated from the midpoints of the geographic bounds in PanTHERIA^52^. For *Tragelaphus angasii*, historical approximate locations recorded in 23 Universal Transverse Mercator coordinates were converted using their zones and hemispheres, assuming WGS84.

Evolutionary distances came from the TimeTree-based phylogeny^53^. We summed the branch lengths connecting each species pair, expressed in millions of years. This is the total evolutionary path separating the species, rather than the age of their most recent common ancestor.

### Accounting for sampling bias

We accounted for uneven study effort in several ways. First, we retained species with at least ten distinct recorded parasite names, reducing the influence of very sparsely documented communities^13^. Species also required complete geographic, evolutionary, taxonomic, and publication-count fields. These criteria yielded 680 species and 30,310 distinct associations. For parasite-type analyses, the ten-name threshold was applied within each type, producing type-specific species sets. Second, we included publication counts as a covariate, following approaches that use research effort to adjust comparative parasite analyses^20,42^. For each host species, we queried PubMed using its scientific name and documented synonyms, including previous genus assignments, combined with OR. We required the title or abstract to contain a host name and at least one of the following terms: parasit*, pathogen*, infect*, disease*, virus, viruses, viral, bacteri*, protozo*, fung*, helminth*, worm*, arthropod*, or ectoparasit*. Asterisks allowed matching alternative word endings. We obtained the number of matching publications through the NCBI ESearch interface, using a single combined query per species so that publications matching multiple synonyms were counted only once. Third, Jaccard similarity expresses sharing relative to the combined community size, reducing the influence of differences in recorded richness compared with raw shared-parasite counts^12,25^. Finally, excluding humans and common domesticated animals reduced the influence of disproportionately studied species^44^.

### Selecting predictors of parasite sharing

We constructed pairwise distance matrices for the host traits (Table S1) and for geographic and evolutionary distance. Numeric trait distances were absolute differences between recorded species-level values, and categorical distances indicated whether species belonged to the same or different categories. Geographic and evolutionary matrices used the distances described above. To assess associations among these predictors, we calculated Spearman Mantel correlations for all 66 unique matrix pairs^54^ (Table S2; Supplementary File S1). Mantel tests were performed using the R package ‘vegan’ version 2.7-6^55^. Each test used species with complete data for the two variables and 9,999 unrestricted permutations of species labels. Two-sided P-values used a plus-one correction. Strong associations included body length with body mass (r = 0.885), body mass with home range (r = 0.568), and body length with home range (r = 0.465). These associations motivated adding only one host trait at a time to the geographic, evolutionary, and study-effort model, avoiding simultaneous inclusion of strongly correlated host traits.

### Models across taxonomic scales

To investigate the importance of geographic distance and evolutionary relatedness for predicting parasite sharing, while controlling for potential covariation among the predictors, we used a generalized linear model for dissimilarity matrices using the generalized dissimilarity modeling (GDM) approach^24^. This extension of matrix regression represents nonlinear increases in dissimilarity using nonnegative spline coefficients and allowed us to infer potential nonlinear functional forms of parasite sharing for each predictor (partial dependence plots). We modeled one minus Jaccard similarity to obtain dissimilarity. Each distance predictor used three spline basis functions, with knots determined from the observed distances, including their median. The model specification was:

(1 – *J*) ∼ η(geographic dissimilarity, evolutionary dissimilarity, publication count dissimilarity)

where the linear predictor η (a scaled combination of interhost species distances based on the selected covariates) was defined by the default GDM formulation, using a linear combination of spline basis functions (analogous to terms in a polynomial regression), as

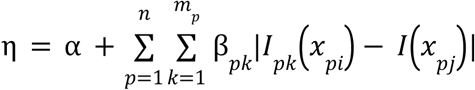

where α denotes the model intercept, *p* is the predictor (*n* in total), *I_pk_* is the k^th^ spline (out of *m_p_* total splines) for variable *x_p_*, and β*_pk_* is the fitted coefficient for *I_pk_*, with the constraint β*_pk_*≥0. By default, a maximum-likelihood estimation approach was used to fit the required coefficients, β*_pk_*. We used the GDM default link function

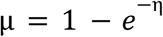

since the response (Jaccard’s dissimilarity index) is a proportion. Geographic, evolutionary, and study-effort contributions were combined additively on the model’s link scale. The GLM variance function (defining how the variance of the predicted response µ depends on the predicted mean) was a modified binomial variance function (the default in GDM) that accounts for proportions (values ranging 0–1)

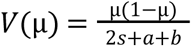

as, by construction, µ = [0, 1] and 2*s* + *a* + *b* is the sum of the total number of parasite species names associated with each host species according to our dataset.

We first fit the GDM model to the whole dataset (full dissimilarity matrices) and recorded the sum of the spline coefficients for each predictor, which measured predictor importance at the class taxonomic level (Mammalia). Next, we subset the dissimilarity matrices to only include the pairwise values (matrix entries) where the species belonged to the same order and fit a GDM model using those to infer predictor importance at the order level. We repeated this step to also infer predictor importance at the family and genus taxonomic levels. The nested analyses included 1,125 same-genus pairs from 449 host species, 9,335 same-family pairs from 656 host species, 41,211 same-order pairs from 679 host species, and all 230,860 pairs from 680 host species.

We measured relative geographic importance as the sum of its fitted spline coefficients divided by the combined coefficient sums for geography and evolutionary distance, and calculated evolutionary importance in the same way. Study effort was excluded from this denominator. The two shares therefore sum to one and describe relative fitted distance contributions, rather than percentages of explained variation.

To account for model uncertainty, we generated 1,000 host-level bootstrap samples. We resampled host species to account for dependence among pairs that include the same host species. Each pair was weighted by multiplying the number of times its two host species appeared in the resample. Intervals were the 2.5th and 97.5th percentiles of retained estimates. Bootstrap comparisons assessed whether the geographic and evolutionary contributions differed at each scale. We checked sensitivity to unequal sample sizes using 1,000 repeated samples of 1,000 host pairs at each taxonomic scale. In an additional equal-host-count analysis, we first selected 449 eligible host species at each scale, then sampled 1,000 pairs and refitted the models. These analyses balanced the sample sizes. All GDM analyses were performed using the R package ‘gdm’ version 1.6.0-7^56^.

### Host traits and recorded parasite host ranges

We added ten PanTHERIA traits separately to the geographic, evolutionary, and study-effort models at each nested scale: body length, body mass, habitat breadth, home range, litter size, maximum longevity, population density, activity cycle, terrestriality, and trophic level^52^ (Table S1). Each analysis used species with the relevant trait available. Numeric traits entered as recorded species-level values. Categorical traits were encoded as the same or different categories. We attempted 1,000 bootstrap samples for each of the 40 trait models and retained finite full-rank fits without replacing failures.

Recorded parasite host ranges were calculated within the retained set of 680 species and summarized by parasite type (Fig. S3). Following a conservative screen of generic parasite names, we calculated the proportion recorded in only one species and the distribution of recorded host-range sizes. These summaries measure documented associations within the study dataset, rather than complete biological host ranges.

### Assessing the importance of taxonomic scale for parasite-sharing predictions

We compared the performance of the two models trained at the taxonomic extremes: genus and class. The genus-trained model and the class-trained model were compared on identical target pairs under whole-genus holdout. For each target comparison, a baseline model assigned every test pair the mean observed sharing among the corresponding training pairs: within-genus training pairs for within-genus targets, and class-level training pairs for class-level targets. The standardized root mean squared error was model error divided by the baseline model error, giving a unitless ratio. Values below one indicated improvement over the baseline. This comparison tests how well a model fitted at one taxonomic scale predicts sharing at another.

We evaluated sharing predictions with five repetitions of five-fold host-species holdout and a separate five-fold holdout of entire genera. Training excluded all pairs containing a held-out species. Performance measures included root mean squared error and mean absolute error. Errors were expressed in percentage points because sharing was measured on a percentage scale. Paired uncertainty comparisons used 2,000 species- or genus-block resamples of evaluation errors, conditional on the fitted models and chosen folds.

### Counterfactual parasite sharing under changing geographic conditions

We used the generalized dissimilarity models and reduced each pair’s range-center distance by 25%, 50%, 75%, or 99%, holding evolutionary distance and study effort fixed. We compared currently observed sharing with predicted sharing under counterfactual scenarios with reduced geographic separation. Observed intervals used 1,000 host-species resamples, and scenario intervals used compatible saved species-bootstrap fits. These scenarios assess conditional model responses, and do not simulate actual range shifts, local contact, or climate-driven transmission. All analyses were conducted in R version 4.6.1^57^.

## Acknowledgments

We are grateful to Sarah Bowden (Centers for Disease Control and Prevention) for early discussions that inspired this project. We also thank Susan Holmes (Department of Statistics, Stanford University), Giulio De Leo (Department of Oceans, Stanford University), Elizabeth Hadly, Dmitri Petrov, and Marcus Feldman (Department of Biology, Stanford University) for continuous discussions and feedback.

## Funding

N.N. was supported by the Philanthropic Educational Organization (PEO) Scholar Award from the International Chapter of the PEO Sisterhood, the Stanford Data Science Scholars program, and the Predoctoral Fellowship from the Stanford Center for Computational, Evolutionary and Human Genomics, and the National Institutes of Health (NIH; R35GM133439). E.A.M. was supported by the National Science Foundation (DEB-2011147, with the Fogarty International Center), the NIH (R35GM133439), the Terman Award, the Stanford King Center for Global Development, the Stanford Woods Institute for the Environment, and the Stanford Center for Innovation in Global Health.

## Author Contributions

N.N. and E.A.M. conceived the study. N.N. collected the data. N.N. performed the analyses. N.N. drafted the manuscript, and all authors contributed to revisions.

## Competing Interest

The authors declare no competing interests.

## Data availability

The data used in this study are publicly available^23,43,45–50,52,53^. Code and formatted data used in this study will be available on Github at the time of publication.

## Notes

### Competing Interest Statement

The authors have declared no competing interest.

