## Supplement for "Ecological opportunity and evolutionary relatedness predict parasite sharing differently across mammalian taxonomic scales"

**This PDF file includes:**

Figures S1 to S4
Tables S1 to S2
Legend for Supplementary File S1

**Other supporting materials for this manuscript include the following:**

Supplementary File S1

### Supplementary Tables and Figures

**Table S1. Morphological and ecological trait variables examined as additional predictors of parasite sharing.** Ranges are the minimum and maximum recorded values among host species in our dataset. Activity cycle, terrestriality and trophic level are categorical variables. Their displayed ranges refer to source category codes, which were modeled as same/different categories rather than ordered numerical distances. Habitat breadth is a count, whereas litter size and population density are species-level numeric summaries that can take fractional values. The data for all variables were obtained from the publicly available PanTHERIA database[52].

| Variable | Description | Unit or datatype | Range |
|---|---|---|---|
| Body length | Average adult length from nose to anus or base of tail | mm | 38.99–4177.03 |
| Body mass | Average adult body mass | g | 2.46–3824539.93 |
| Habitat breadth | Number of habitat layers occupied: fossorial, ground-dwelling, arboreal or aquatic | Integer count | 1–3 |
| Home range size | Average home-range area utilized by the species | $km^2$ | 0.000020373–2771.22 |
| Litter size | Average number of offspring per litter | Numeric mean (offspring/litter) | 0.84–11.23 |
| Maximum longevity | Maximum recorded lifespan | Months | 11–960 |
| Population density | Average number of individuals per unit area | Individuals/$km^2$ (numeric) | 0.004822132–57067.85 |
| Activity cycle | Daily activity pattern | Categorical: 1) nocturnal only; 2) mixed or crepuscular activity; 3) diurnal only | Codes 1–3 |
| Terrestriality | Classification of terrestrial versus other habitat use | Categorical: 1) fossorial and/or ground-dwelling only; 2) above-ground dwelling | Codes 1–2 |
| Trophic level | Diet-based trophic classification | Categorical: 1) herbivore; 2) omnivore; 3) carnivore | Codes 1–3 |

**Table S2. The most collinear variables in the dataset.** These are Mantel test results to test for collinearity between dissimilarity matrices of model predictors for parasite sharing. All variable combinations and their collinearity results are included in the Supplementary File S1.

| **Variable 1** | **Variable 2** | **Spearman correlation coefficient (r)** | ***P*-value** |
|---|---|---|---|
| Body length | Body mass | 0.885 | 0.0001 |
| Body mass | Home range size | 0.568 | 0.0001 |
| Body length | Home range size | 0.465 | 0.0001 |
| Population density | Litter size | 0.362 | 0.0001 |
| Activity cycle | Phylogenetic distance | 0.229 | 0.0001 |
| Body mass | Population density | -0.293 | 0.0001 |
| Population density | Home range size | -0.322 | 0.0001 |

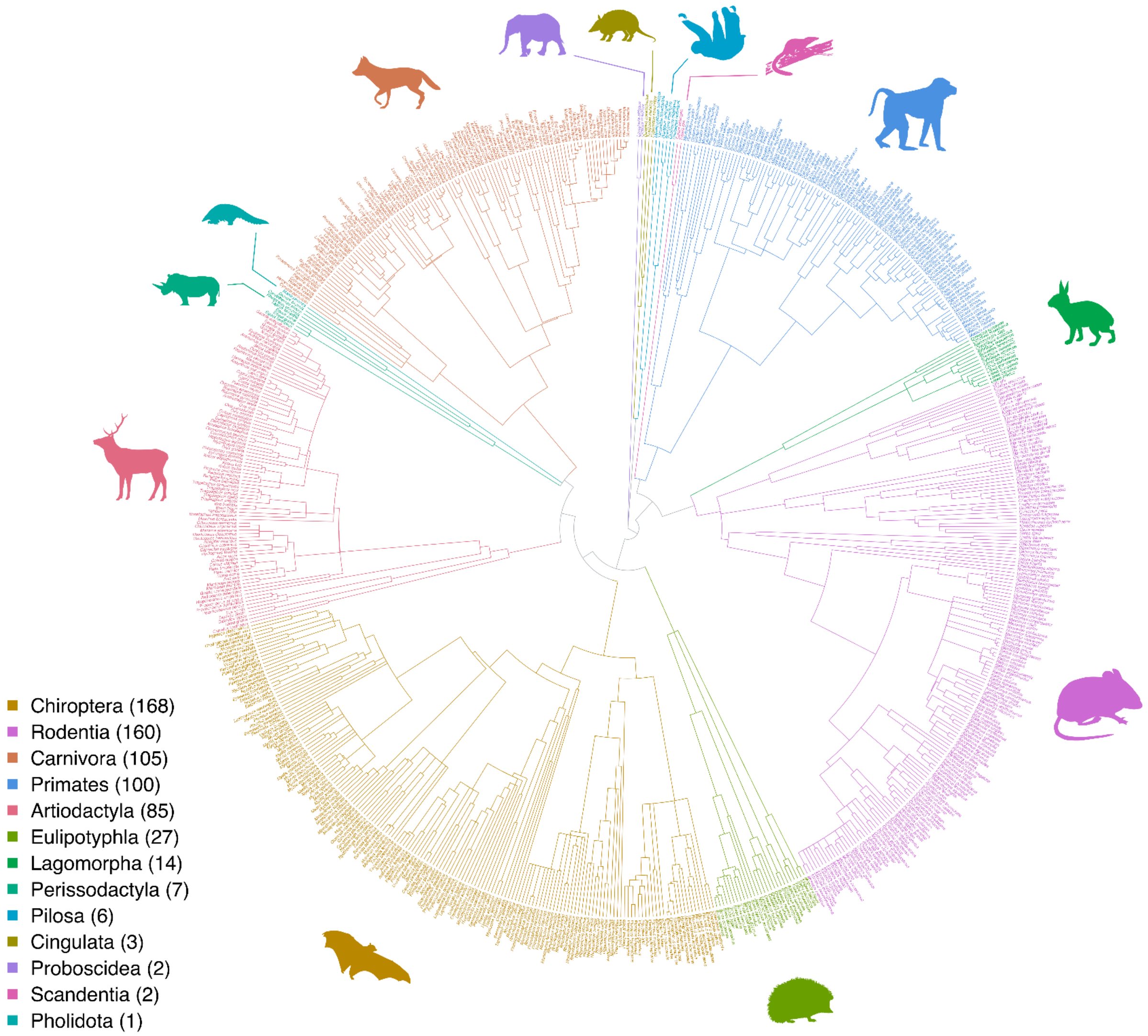


**Fig. S1. Evolutionary relationships among the 680 mammalian host species included in the parasite-sharing analyses.** The circular phylogeny was pruned from the TimeTree phylogeny used to calculate host evolutionary distances. Branch lengths represent estimated evolutionary time. Species names and branches within each of the 13 orders are colored consistently with the host-order parasite-sharing network (Fig. 1). Branches connecting different orders are gray. Numbers in the legend indicate the number of host species in each order. Silhouette icons were obtained from phylopic.org.

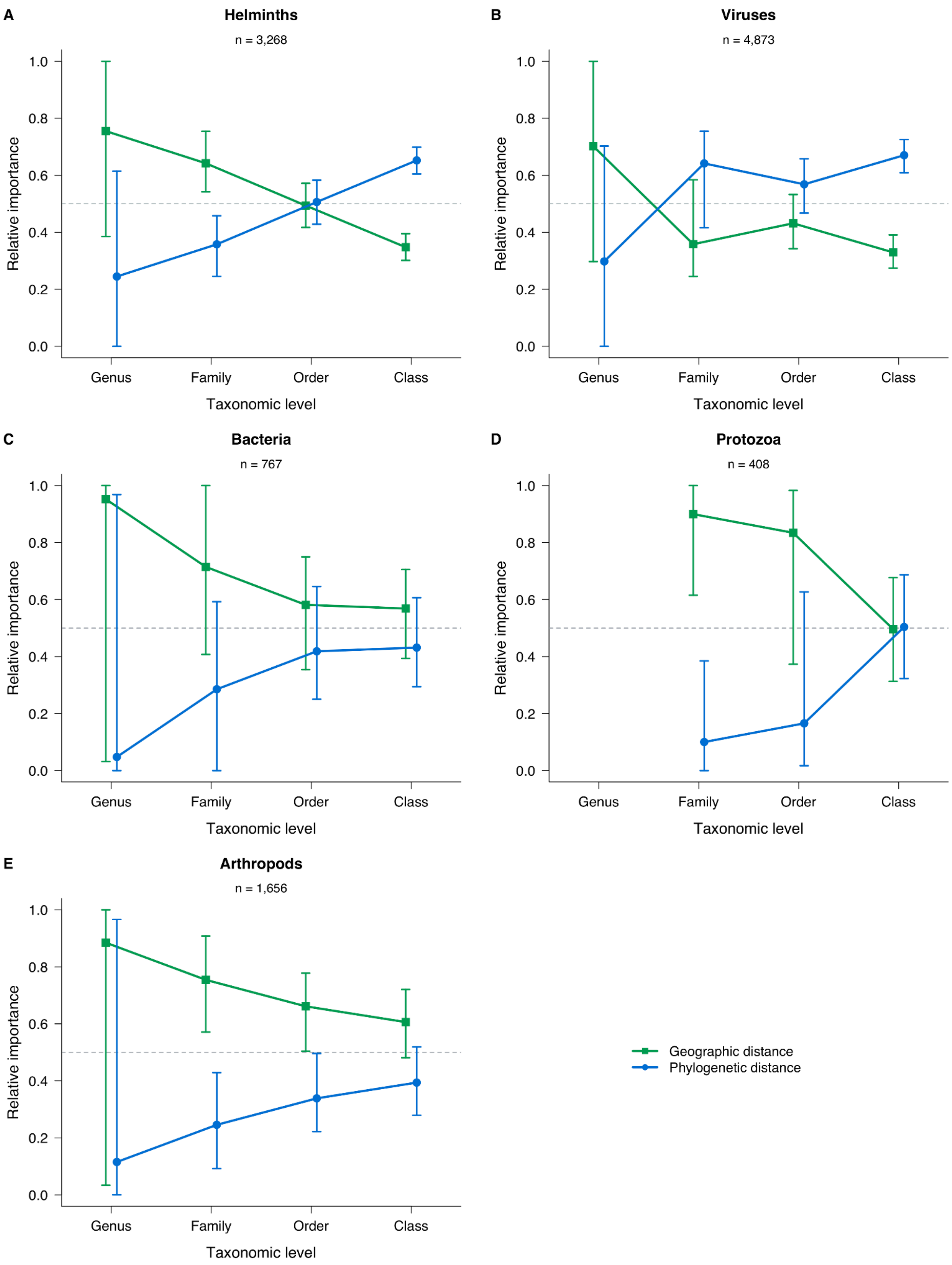
A
Helminths
n = 3,268
B
Viruses
n = 4,873
C
Bacteria
n = 767
D
Protozoa
n = 408
E
Arthropods
n = 1,656
Relative importance
1.0
0.8
0.6
0.4
0.2
0.0
Genus
Family
Order
Class
Taxonomic level
Geographic distance
Phylogenetic distance

**Fig. S2. Relative importance of geographic and evolutionary distance for parasite sharing across taxonomic scales by parasite type.** Panels show models fitted separately for (A) helminths, (B) viruses, (C) bacteria, (D) protozoa and (E) arthropods ("n" indicates the number of parasite species included for each parasite category). No genus-level protozoan model was available due to lack of data. Green squares represent geographic distance between host range centers, and blue circles represent phylogenetic distance. Points show each predictor's summed spline coefficients divided by the combined coefficient sums for geography and phylogeny, so their relative contributions sum to one within each taxonomic level. Lines connect estimates across models fitted to pairs within the same genus, family, order or class. Vertical bars show 95% percentile intervals from finite host-bootstrap fits, with 1,000 attempted replicates per model. Relative importance is calculated separately within each draw. The dashed line marks equal contributions.

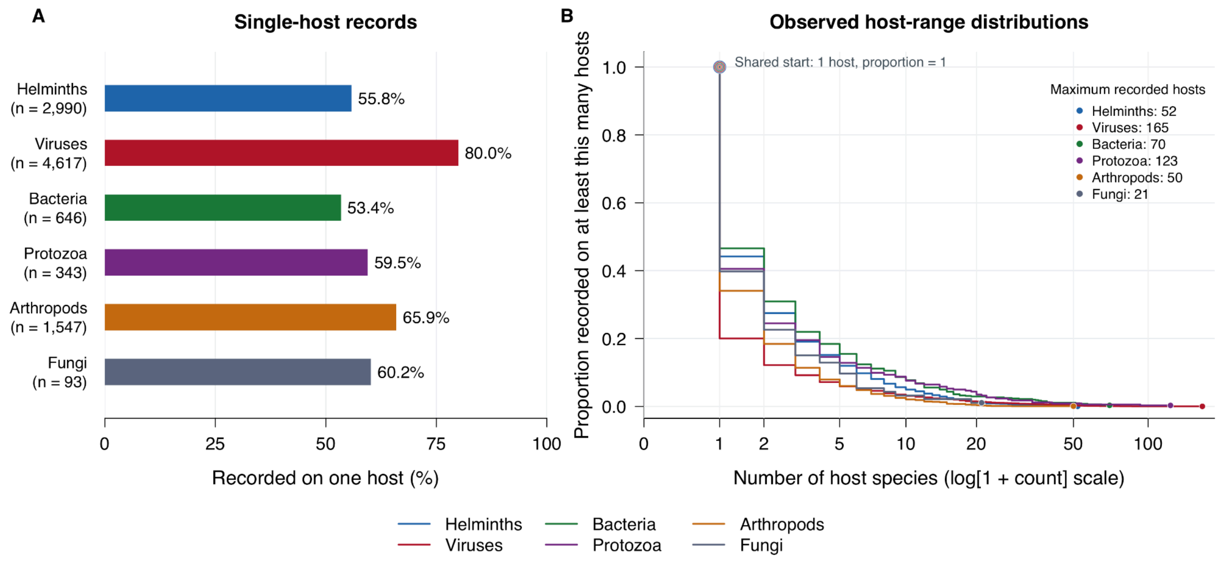


**Fig. S3. Recorded parasite host ranges in the dataset.** Host ranges were calculated from recorded associations within the same set of 680 mammalian host species. (A) Percentage of retained parasite labels recorded on only one host species, by parasite type. Numbers in parentheses indicate the number of retained parasite labels, and values beside bars give the corresponding percentages. (B) Proportion of retained parasite labels recorded on at least the number of host species indicated on the x-axis, displayed on a log(1 + count) scale. Each curve begins at one host with a proportion of one. Terminal points and the inset legend identify the maximum recorded host range for each parasite type. Colors identify parasite types consistently across panels.

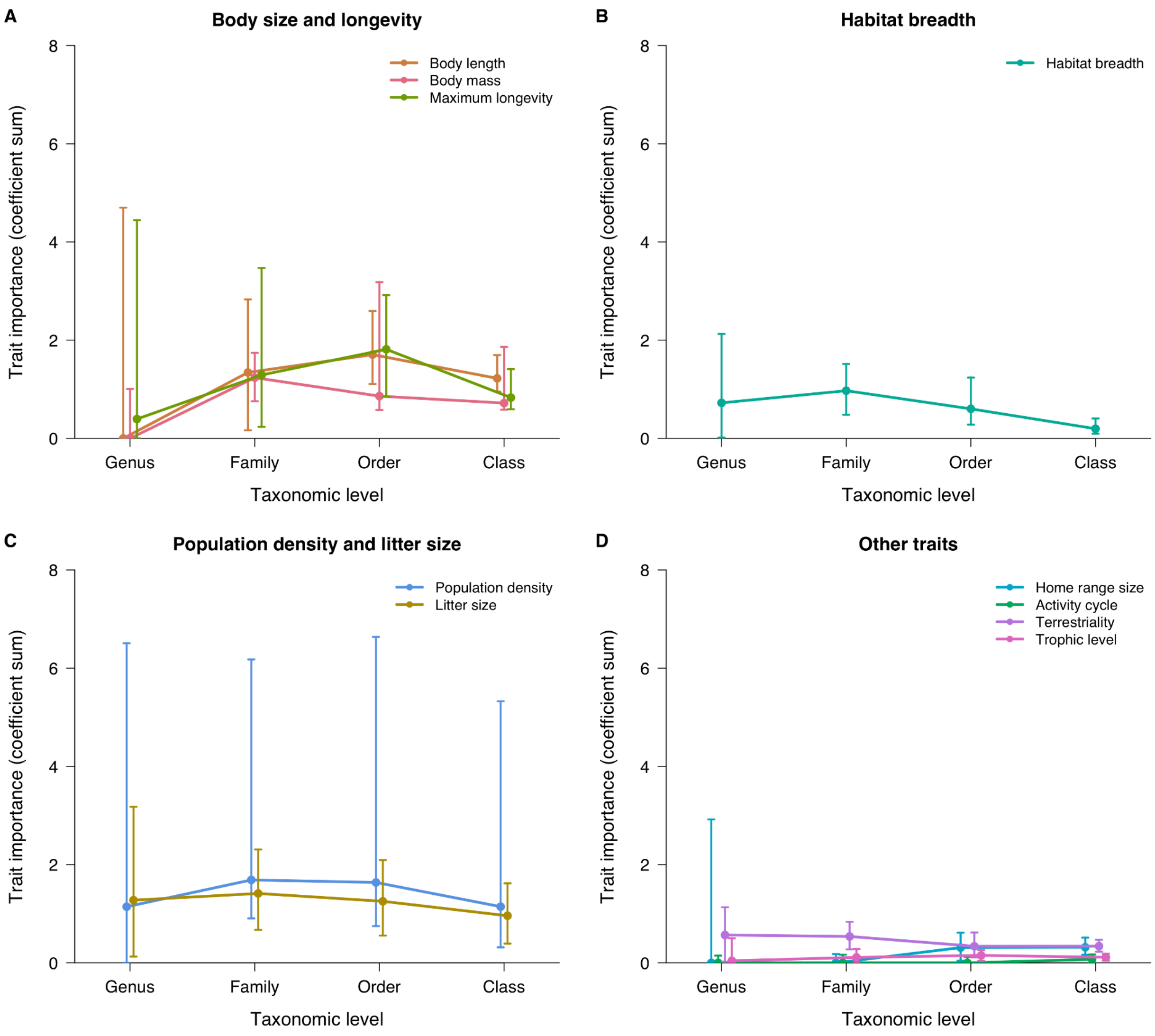


**Fig. S4. Contributions of individual host traits to parasite community dissimilarity across mammalian taxonomic scales.** Each trait is added separately to a generalized dissimilarity model containing geographic distance, phylogenetic distance and study effort. Points show summed trait spline coefficients, and lines connect estimates for same-genus, same-family, same-order and all-mammal pairs. Error bars show pointwise 95% percentile intervals from 1,000 host bootstrap fits.

**Supplementary File S1. Pairwise associations among host predictor distance matrices.** Spearman Mantel correlation coefficients (r) and two-sided permutation P-values are shown for all 66 unique pairs of ten host traits, geographic distance and phylogenetic distance, ordered from highest to lowest signed r. Numeric trait distances are absolute differences in recorded host values, where activity cycle, terrestriality and trophic level use same/different-category distances. Each comparison uses pairwise complete hosts from the 680-host analysis universe. Sample size denotes the number of host species with both predictors available. P-values use 9,999 host-label permutations with a plus-one correction (the minimum attainable P-value is 0.0001).
